# Mapping Lung Cancer Ventilation Dynamics: A Pilot Mouse Study Using Functional Imaging and Lung Mechanics

**DOI:** 10.1101/2025.07.06.661887

**Authors:** Ronan Smith, Nicole Reyne, Daniel Batey, Nina Eikelis, Marie-Liesse Asselin Labat, Martin Donnelley

**Affiliations:** Robinson Research Institute, University of Adelaide, South Australia; Adelaide Medical School, University of Adelaide, South Australia; Respiratory and Sleep Medicine, Women’s and Children’s Hospital, South Australia; Personalised Oncology Division, Walter and Eliza Hall Institute of Medical Research, Melbourne, Australia; Department of Medical Biology, The university of Melbourne; 4DMedical, Victoria, Australia

**Author notes:** The authors contributed equally to the work (Ronan Smith, Nicole Reyne).

**Keywords:** Lung cancer, animal models, lung disease, lung function, X-ray Velocimetry, Permetium, flexiVent

## Abstract

*In vivo* models that replicate and reproduce human lung cancer and its response to therapy are necessary for the development of new therapeutic strategies and understanding drug resistance. Imaging lung tumors in live animals to monitor tumor growth and response to therapy is challenging due to the location of the lungs and their constant movement during breathing. Additionally, methods such as computed tomography (CT) only provide structural information and not functional information about how well the lungs are working.

X-ray velocimetry (XV) is a novel functional lung imaging technique that generates 3D maps of regional lung expansion during breathing. In other lung diseases it has been shown to provide spatial information on where ventilation changes occur. The aim of this pilot study was to use XV and flexiVent lung mechanics assessments to determine the effect of tumor growth on lung function in mice at 2- or 3-weeks post tumor induction, and to evaluate the efficacy of these two tools.

Histological analysis showed that tumour growth was not uniform between animals. At 3-weeks post tumor induction, some XV ventilation and flexiVent lung mechanics parameters were significantly different from baseline metrics. In addition, the forced expiratory volume, small-scale ventilation heterogeneity, and the average CT gray value correlated with the tumour counts from the histology. In some mice XV revealed localised regions with altered expansion rates.

This pilot study demonstrated that changes in lung function can be identified following tumor induction, and that the model and techniques could be used in the future to determine response to anti-tumor drugs.

## Introduction

Lung cancer remains the leading cause of cancer-related deaths worldwide, with a five-year survival rate of less than 20% [1], primarily due to late diagnosis and limited treatment options. Non-small cell lung cancer (NSCLC), which accounts for approximately 85% of lung cancer cases, frequently harbours genetic mutations that drive tumor growth and progression [2]. In lung adenocarcinoma (LUAD), the most common subtype of NSCLC, the most common oncogenic driver mutations are mutations in *KRAS* (∼30% of LUAD cases) and loss of function mutations in the tumour suppressor gene *TP53* (∼50% of LUAD cases [3]). While activation of the *KRAS* oncogene in murine lungs induces lung adenoma formation [4], co-occurring *TP53* mutation accelerates tumour onset and the invasive properties of the tumours leading to lung adenocarcinoma formation [5].

Mouse models that recapitulate the genetic and pathological features of human lung adenocarcinoma have been extensively used to study lung cancer initiation, progression and response to therapy. Cell lines established from the *KRAS*^*G12D*^*/TP53*^*mut*^ animal models [6] further allow for the establishment of lung tumors in a controlled and reproducible manner, enabling tumor growth dynamics, immune responses, and potential therapeutic interventions to be studied *in vivo*. One of the major challenges in the use of animal models of lung cancer is to be able to monitor tumor burden and its impact on respiratory function in live animals.

Current methods to assess tumour burden *in vivo* in pre-clinical models utilise X-ray computed tomography (microCT) [7], or require the use of luciferase-expressing cells for bioluminescence imaging [8]. While microCT provides an accurate overview of tumor location, distinguishing tumour from normal tissue can be difficult without contrast agents. On the other hand, bioluminescence can be sensitive and easy to use, however signal attenuation varies depending on how deep the tumors are in the thorax, reducing quantitative accuracy and precise information on location. In addition, both methods do not provide information on the impact of tumour burden on lung health and respiratory function.

The flexiVent system is the current gold standard for assessing lung function in small animal models. The flexiVent applies the forced oscillation technique to provide precise measurements of airway resistance, lung compliance, and elastance, offering detailed insights into how tumors alter pulmonary mechanics. In addition, negative pressure forced expiration (NPFE) testing can be used to assess airflow in a manner analogous to human spirometry. As lung tumors develop, they would be expected to cause airway obstruction, reduced compliance, impaired gas exchange and altered airflow, which the flexiVent should be able to quantify. To date there have been no reports of the flexiVent being used to characterise changes in lung mechanics from lung cancer.

X-ray Velocimetry (XV) functional lung imaging has emerged as a powerful, non-invasive technique for assessing regional ventilation in preclinical models [9-13]. XV imaging allows real-time visualization of lung motion and airflow distribution, capturing subtle changes in regional lung mechanics and ventilation heterogeneity. There are no reports of XV functional lung imaging being applied to lung cancer. However, XV could be particularly valuable for evaluating how lung tumor development affects localized airflow and how interventions may restore normal ventilation patterns. Unlike traditional lung function tests, which provide global measurements, XV imaging provides regional insights into tissue motion and lung ventilation, potentially enabling a more comprehensive understanding of tumor-induced lung dysfunction.

This pilot study examined a previously established mouse model of lung cancer [6] using flexiVent lung function testing and XV imaging. The aims were to begin to assess the timing of lung cancer progression, and to evaluate the efficacy of these tools in a cancer model. We hypothesised that these methods could provide critical insights into tumor biology and lung mechanics.

## Methods

### Animals

All animal procedures were approved by the Walter and Eliza Hall Institute (2023.006) and South Australian Health and Medical Research Institute (SAM-23-065) Animal Ethics Committees. The parts of the ARRIVE guidelines that pertained to a pilot study were adhered to [14]. All experiments were conducted in accordance with the National Health and Medical Research Council Australian Code of Practice of the Care and Use of Animals for Scientific Purposes.

Lung cancer-bearing mice were supplied by the Walter and Eliza Hall Institute, and were generated by injection of a tumor cell line derived from a murine lung tumor that grew in a cre-inducible *Kras*^*G12D*^;p53^Δ/Δ^ C57BL/6 mouse model. Female age-matched mice received intravenous tail-vein injection of 1 × 10^5^ *Kras*^*G12D*^;p53^Δ/Δ^ lung cancer cells into C57BL/6 mice two or three weeks prior to the imaging, as previously described [6]. Mice were housed in a conventional facility with a 12-h light/dark cycle and food and water were provided *ad libitum*.

### X-ray Velocimetry (XV) imaging

All XV imaging was performed as described previously [9, 12]. Mice were anaesthetised with an intraperitoneal injection of a mix of 1 mg/kg medetomidine (Ilium, Australia) and 75 mg/kg of ketamine (Ceva, Australia). Once anaesthetised the mice were tracheostomised and a short endotracheal tube (ET; 18 Ga BD Insyte plastic cannula bevel-cut to 15 mm length) was inserted. Mice were then secured head-high in a holder and placed onto the translation/rotation stage of a Permetium preclinical XV scanner (4DMedical, Australia) [10], configured with a sample to detector distance of 1700 mm. Mice were connected to a pressure-controlled small animal ventilator (4DMedical Accuvent 200, Australia) and ventilated at a 150 breaths/min (200 ms inspiration and 200 ms expiration; I:E ratio of 1:1) with a peak inspiratory pressure of 14 cmH_2_O, and positive end-expiratory pressure of 2 cmH_2_O. An XV scan was acquired at a framerate of 25 Hz, with 10 phase-points per breath and 600 projections per phase-point. The X-rays were generated by a Rigaku MM09 x-ray source (40 kV, 2 mA) with a 40 μm molybdenum filter, and detected by a Varex 2020DX flat-panel detector (1024 × 1024 pixels), with geometric magnification giving an effective pixel size of 25.7 μm in the sample plane.

Specific ventilation volumetric data was produced by 4DMedical using their proprietary three-dimensional cross-correlation algorithm to quantify tissue displacement throughout the breath. Specific ventilation is defined as the change in volume of a voxel of the lung, divided by the volume of the voxel at the start of the breath. Unless explicitly stated, in this manuscript any reference to XV refers to specific ventilation measured between peak exhalation and inspiration. XV ventilation maps can be viewed by taking 2D slices in any plane. The slices shown here have anatomical features such as bone from the CT overlaid onto the ventilation data. Regions of interest in specific ventilation maps can be segmented in the same way as regions in CT, as we have demonstrated previously [15].

From the specific ventilation volume, mean specific ventilation (MSV), ventilation defect percentage (VDP; the percentage of the lung with the specific ventilation below a threshold, typically 60% of the MSV), ventilation heterogeneity (VH; the interquartile range divided by the mean, which gives a measure of the ventilation variance across the lung) and tidal volume (V_T_) were calculated. For this study we report the normalised ventilation defect percentage (nVDP), which was calculated using 60% of the MSV of the control population to define a threshold for defective regions, as described by Reyne et al. [11, 12]. VH was split into small-scale (VH_SS_) and large-scale (VH_LS_), showing heterogeneity across small (i.e. intra-lobe) and large (i.e. inter-lobe) spatial scales, respectively. These were calculated by applying high-pass and low-pass spatial filters before calculating VH.

### flexiVent respiratory mechanics

After XV scan acquisition, lung function assessments were performed using a flexiVent FX small animal ventilator (SCIREQ, Montreal, Canada) as previously described [12]. To suppress spontaneous breathing during lung function measurements, the mice received an intramuscular dose of 0.6 mg/kg vecuronium bromide after being connected to the flexiVent. Briefly, the flexiVent was fitted with a FX2 mouse module and NPFE forced expiration extension, and operated by flexiWare v8.0 software. Mice were connected to the flexiVent via the 18 Ga cannula, ventilated at a respiratory rate of 150 breaths/min, inspiratory to expiratory ratio of 2:3, tidal volume 10 mL/kg, and positive end expiratory pressure (PEEP) of 3 cmH_2_O. Lung mechanics measurements were made using a mouse mechanics scan script that was repeated three times – consisting of Deep Inflation (IC inspiratory capacity), SnapShot-150 (R_rs_ respiratory system resistance; C_rs_ respiratory system compliance), Quick-Prime 3 (R_n_ Newtonian resistance; G tissue damping; H tissue elastance), PVs-P (C_st_ static compliance; K curvature; Area) and NPFE (FEV0.05 forced expiratory volume 0.05 seconds; FVC forced vital capacity; FEF0.05 forced expiratory flow 0.05 seconds) perturbations – with the Scireq automated algorithms set to default values [12, 16]. For each parameter three measurements were made per mouse and these were averaged. Data was excluded if the coefficient of determination was less than 0.9 for each model. After lung mechanics were assessed, the mice were humanely killed with an intraperitoneal injection of sodium pentobarbitone (∼200 mg/kg).

### Computed tomography

A computed tomography (CT) volume can be reconstructed from the data acquired during the XV scan (referred to as the breathing CT), however due to respiratory and cardiac motion, the sharpness of this scan is limited. As the breathing CT and XV volumes align, the same segmentation was applied to the breathing CT. From this, the average CT gray values across the lung were calculated for each animal. To produce a sharper image of the lung structure, a second CT scan was performed after the flexiVent testing was performed, the animal was humanely killed and the heart had stopped beating (referred to as the post-mortem CT). That scan used 3000 projections, with all other Permetium parameters the same as in the XV scan described previously. The voxel size in both the reconstructed volumes was 25.7 μm.

### Histological analysis

The lungs were instillation-fixed *in situ* under gravity at a constant fluid pressure of 25 cmH_2_O with 10% neutral buffered formalin (NBF). The lungs and heart were removed and suspended in 10% NBF overnight and then transferred into ethanol. The lungs were then embedded in paraffin. Tissue sections (5 μm) were cut from the formalin-fixed paraffin-embedded left lung blocks, and stained with Haematoxylin and Eosin. All slides were imaged in brightfield with an Olympus VS200 slide scanner, and images analysed with QuPath software [17]. Cell segmentation and cell counting was performed in an automated manner using the QuPath Cell Detection algorithm applied to full-resolution images. Tumor burden was quantified in QuPath using Artificial Neural Network Multilayer Perceptron (ANN_MLP) object classifiers trained on manually annotated tumour and lung tissues.

### Statistics

All statistical analyses were performed in R version 4.4.1 [18]. Separate t-tests were used to determine whether the weight of the control and cancer mice was significantly different. For the XV and flexiVent analyses the statistical findings were expressed in terms of estimated marginal means and confidence intervals returned from linear models fitted to the data. For every flexiVent and XV parameter, a standard linear regression model was fitted using the “lm” function with a fixed effect of treatment. Post hoc pairwise comparisons for the fitted models were carried out using the “emmeans’’ package [19]. The tumor count data was log_10_ transformed, the same linear regression model applied, and the estimates and confidence intervals assessed on the log scale. Due to the small sample size the results are graphically represented as individual data points and boxplots (median, box represents the IQR and the whiskers 1.5*IQR) to show variability, and the relationships between the ventilation maps and parameters. The Pearson correlation coefficient was calculated to measure the relationship strength between the log_10_ transformed tumor count data and each of the flexiVent and XV parameters, with results reported when *p*<0.05.

## Results

Data was collected from control (n=2) mice, and lung-tumour bearing mice 2-weeks (n=3) or 3-weeks (n=3) after tumour cell injection. The mice were the same age, and there was no significant difference in the weight of the three groups. All mice tolerated the tumor induction and XV imaging. Lung mechanics could not be assessed by flexiVent in one of the 3-week animals (referred to as C1) due to a system leak.

### X-ray Velocimetry imaging

XV produces a three-dimensional map of the specific ventilation at around 6000 voxels (0.4 mm x 0.4 mm x 0.4 mm in size) across the lung. The 3-week group exhibited much more obvious reductions in specific ventilation (red regions) across the entire lung than the control or 2-week groups (Figure 1a-c). VH_LS_ was significantly reduced in the 3-week group compared to control, but there were no statistically significant differences in the mean specific ventilation (MSV), normalised ventilation defect percentage (nVDP), tidal volume (V_T_) or ventilation heterogeneity (VH) between the control and tumor-bearing mice at either two or three weeks post tumor induction (Figure 1d-i).

**Figure 1:**
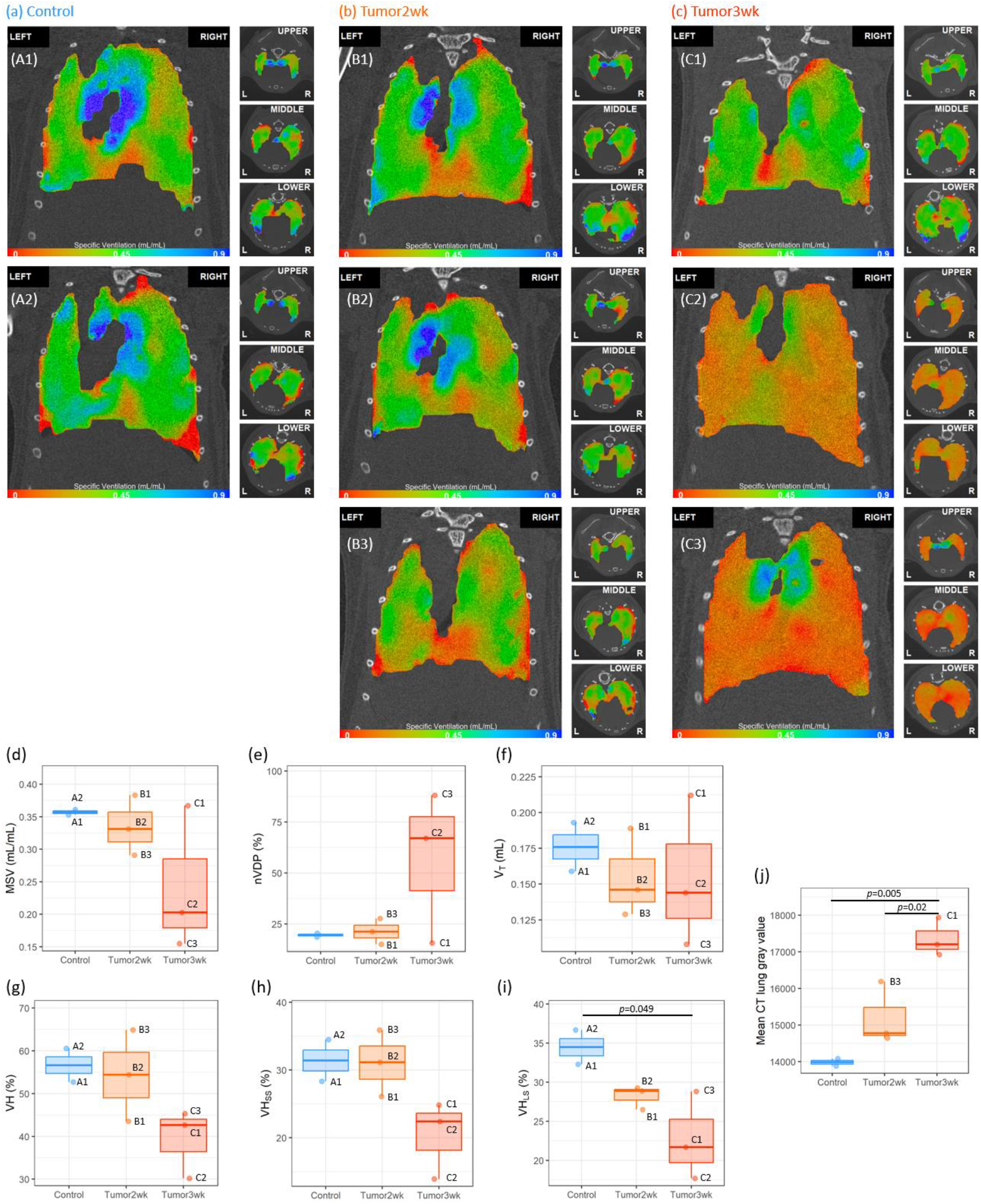
XV ventilation maps. Coronal and axial slices from the 3D ventilation maps produced by XV from (a) control mice, (b) the 2-week group, and (c) the 3-week group. Green indicates average ventilation, red below average ventilation and blue above average ventilation. (d-i) Global XV parameters were variable, but only the large-scale ventilation heterogeneity (VH_LS_) was significantly lower in the 3-week group compared to control. (j) The mean CT gray value was significantly lower in the 3-three week group compared to control. Note that the individual mice are labelled to enable their characteristics to be tracked across the datasets.

Further inspection of the ventilation maps showed that some of the mice had small, localised regions of low ventilation (Figure 2). These regions could be seen in two of the 2-week mice (mice B2 and B3 in Figure 2a and 2b), with no obvious structural changes visible in these areas in the post-mortem CT scan (Figure 2d). Similar localised low ventilation regions could also be seen in a 3-week mouse (C1 in Figure 2c) that did not have the large clear low ventilation regions seen in Figure 1.

**Figure 2:**
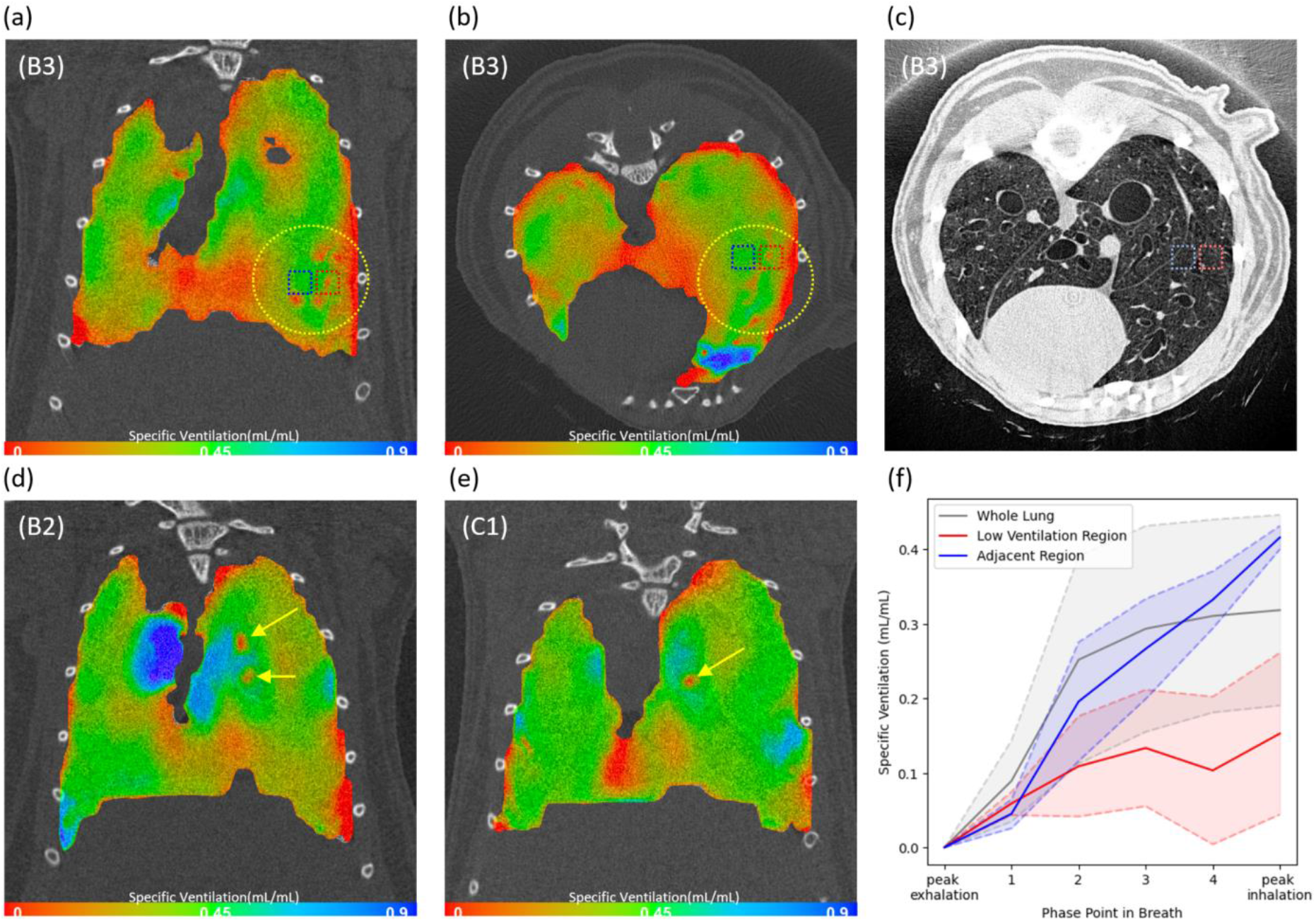
Case studies of XV ventilation maps showing localised areas of very low ventilation. (a-b) Multiple small regions of low ventilation (yellow circles) were seen in the coronal and axial slices from mouse B3 from the 2-week group, although (c) no obvious change was visible in the post-mortem CT slice from the corresponding area (note that the lungs moved between the XV and post-mortem CT scans because the animal was removed from the scanner for flexiVent testing, so the images do not correlate perfectly). (d-e) Similar regions (yellow arrows) were identified in mouse B2 from the 2-week group and in the least severely affected mouse C1 from the 3-week group. Green indicates average ventilation, red below average ventilation and blue above average ventilation. (f) Specific ventilation across the five phase points throughout the breath for mouse B3 shown in a-c. The grey line is the inflation of the whole lung, the red line is the inflation in one of the areas of interest (red box in a-b), and the blue line is the inflation in a nearby region of normal ventilation (blue box in a-b). The shaded areas are the standard deviations in the respective regions.

**Figure 3:**
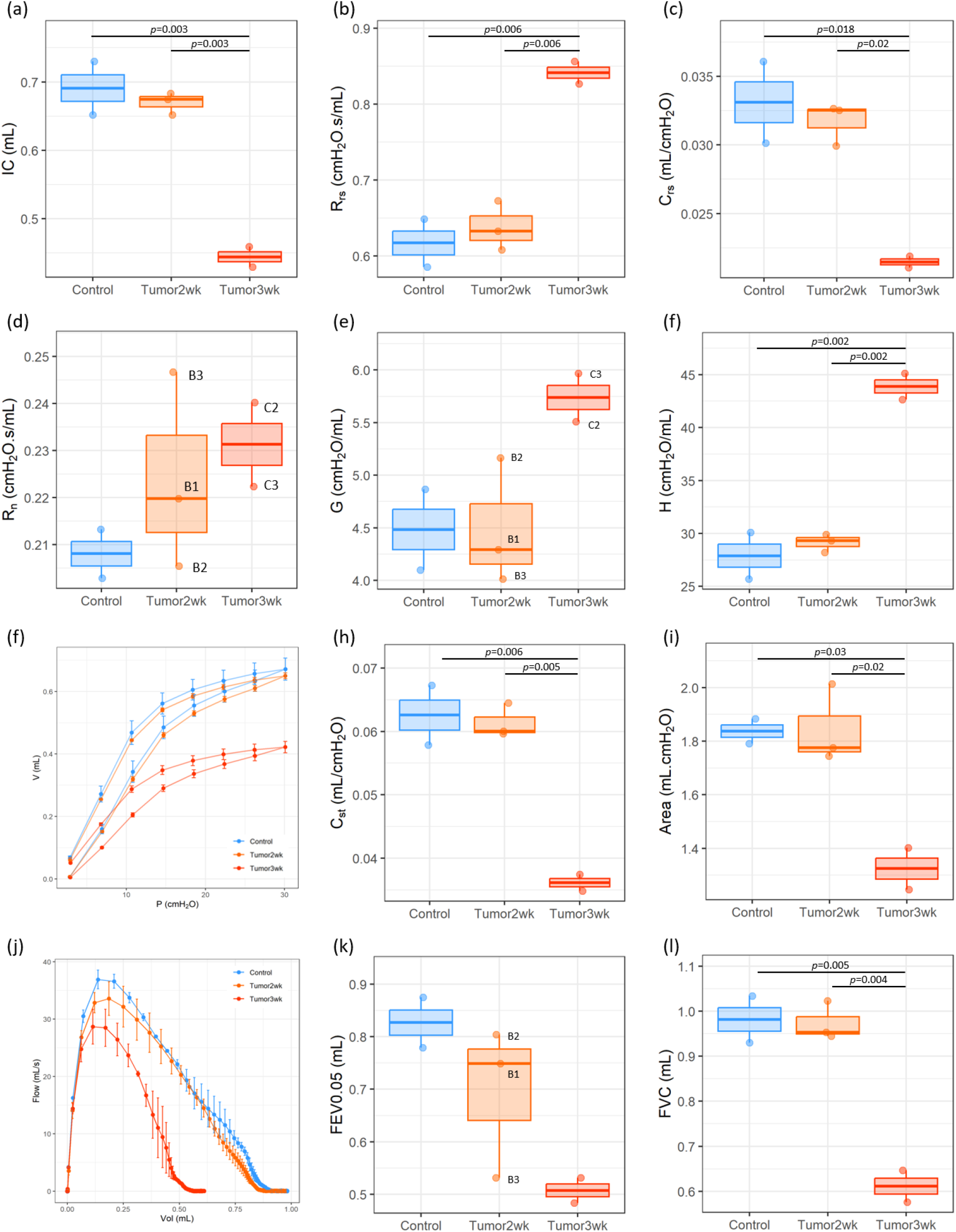
Lung mechanics results. The mechanics of the respiratory system are significantly altered at 3-weeks, but not 2-weeks post-tumor induction. (a) Inspiratory capacity from the deep inflation manoeuvre. (b-c) Respiratory system resistance and compliance from the single compartment model. (d-f) Central airway resistance, tissue elastance, and tissue damping from the forced oscillation technique. (g-i) The stepwise pressure-volume loop, static compliance, and pressure-volume loop area. (j-l) Flow-volume plot and flow parameters from the negative pressure forced expiratory test.

Further analysis was performed to investigate these small localised ventilation defects. We defined a region of interest of approximately 0.8 × 0.8 × 0.8 mm surrounding one of these regions (the position is shown at the yellow box in Figure 2a). The MSV during the breath in this region of interest was compared to an adjacent identically sized region, to explore any temporal changes. Curves showing the expansion of these regions throughout inspiration compared to the expansion of the whole lung are shown in Figure 2e, with the shaded areas showing the standard deviation of specific ventilation within these regions. The inflation of the whole-lung was non-linear, with the lung inflating much more rapidly at the beginning of the breath, before plateauing. A similar pattern to this was seen in the region adjacent to the region of interest. However, within the region of interest, the inflation was very different, slowly expanding across the breath and remaining much lower than the rest of the lung.

The CT scans showed that the tumors were more dense than the normal lung tissue, with the mean CT gray level increasing significantly at the 3-week time point (Figure 1j).

### flexiVent respiratory mechanics

Selected results from the flexiVent testing are shown in Figure 3. There were no significant differences between the control and 2-week tumor groups for any of the tests. However, at 3-weeks, mice exhibited significant reductions in inspiratory capacity (IC) and respiratory system compliance (C_rs_), along with an increase in respiratory system resistance (R_rs_). Tissue elastance (E_rs_) was significantly increased at 3-weeks. Average pressure volume loops constructed from mean data showed a downward shift in the pressure-volume relationship of the three week mice, with corresponding reductions in static compliance (C_st_) and PV loop area. The negative pressure forced expiratory test showed altered flow characteristics and a reduced forced vital capacity (FVC) at 3-weeks.

**Figure 3:**
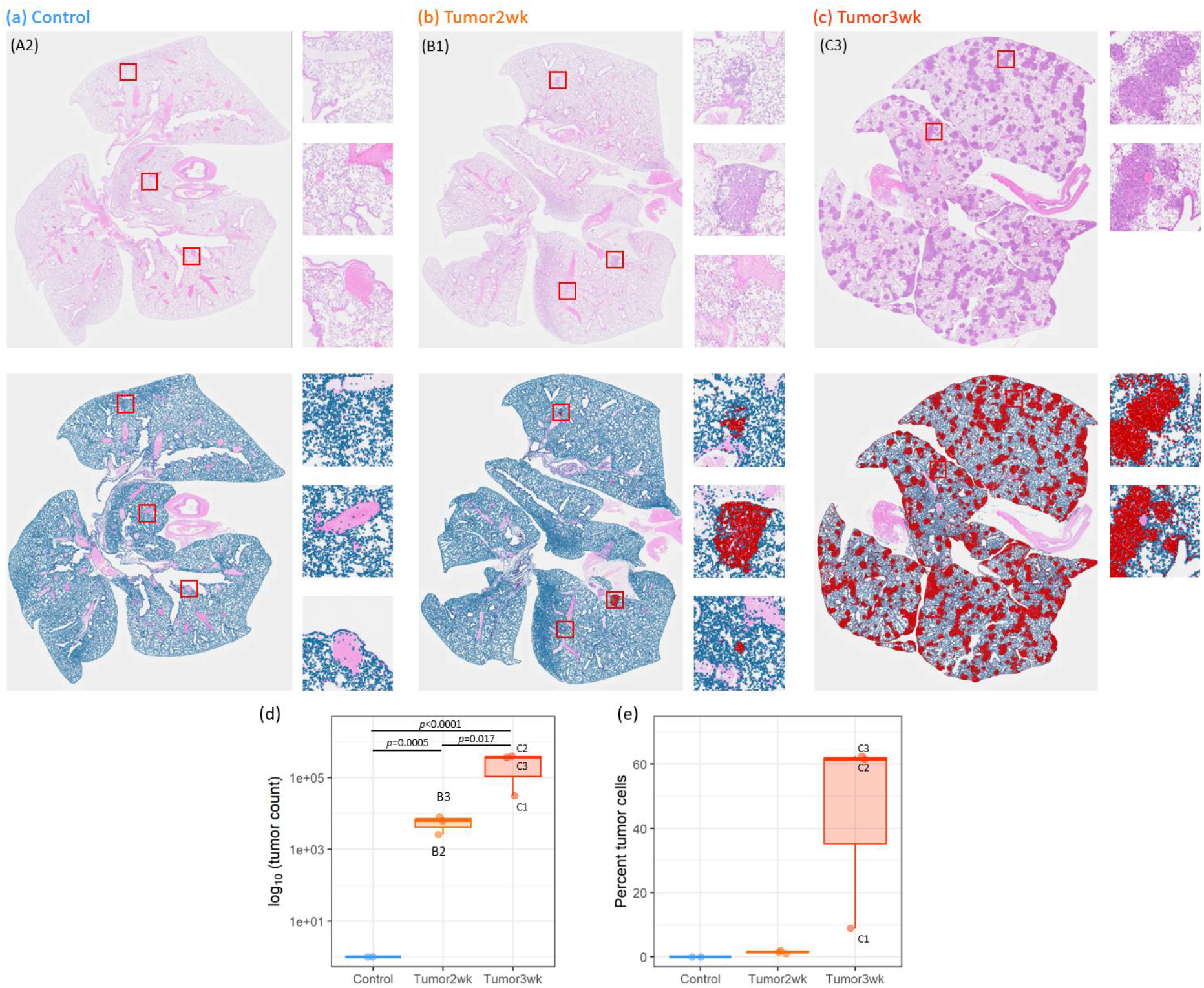
Histological images of tumor-containing lung tissue. Representative images from (a) Control, (b) 2-week and (c) 3-week tumor mice. Tumors were quantified and represented as (d) total tumor counts and (e) tumor burden as a percent of total lung area.

### Lung Tissue Histology

Histological analysis was performed on the excised lung tissue. Tissue sections from both control animals exhibited normal airway and alveolar structure. In contrast, in the two lung cancer groups obvious tumors were identified across the lung, with tumor burden representing 1.5% of the lung area in the 2-week group and 44.3% in the 3-week group (Figure 3). Mouse C1, which appeared to have relatively normal mean specific ventilation (MSV), normalised ventilation defect percentage (nVDP), tidal volume (V_T_) and ventilation heterogeneity (VH) as measured by XV (Figure 1), was the mouse that had the lowest tumour burden by histology. Correlations between log_10_(tumor count) and each of the XV and flexiVent parameters were assessed, with the relationships to FEV0.05, VH_LS_ and mean CT gray value found to be statistically significant (Figure 4).

**Figure 4:**
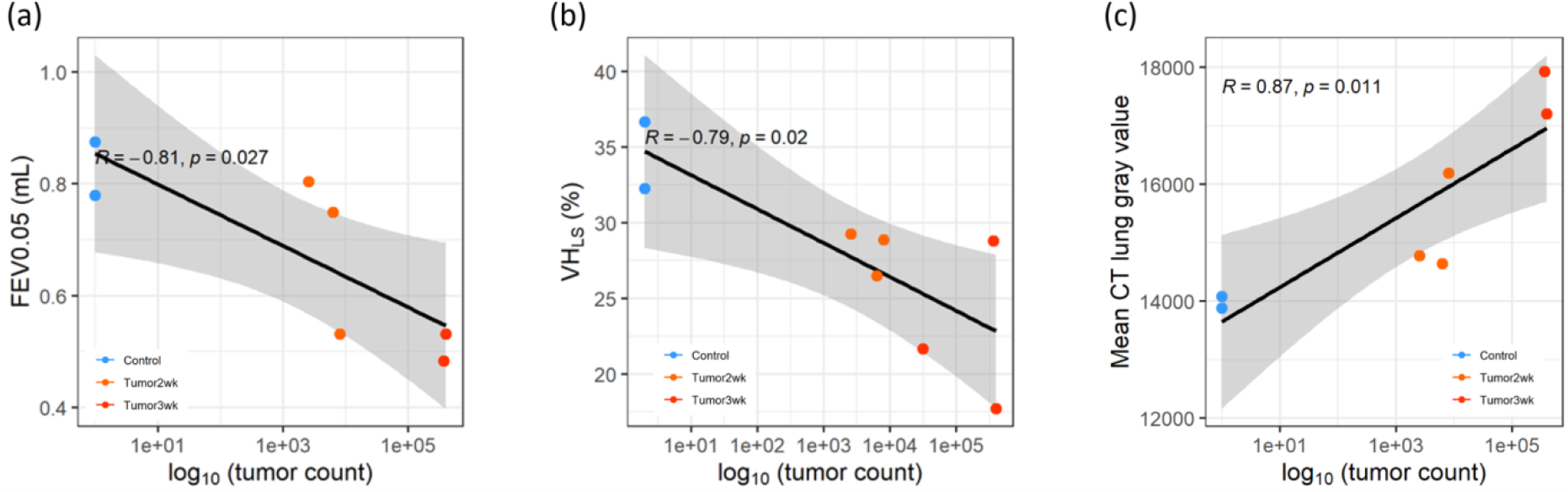
Statistically significant correlations between tumor burden and the (a) flexiVent parameter FEV, (b) XV parameter VH_LS_, and (c) average CT gray value. There were no other significant correlations between the tumor burden and any other parameter.

## Discussion

The aim of this pilot study was to evaluate a previously established mouse model of lung cancer using flexiVent lung function testing and X-ray Velocimetry imaging to assess the timing and extent of lung cancer progression. XV ventilation maps (Figure 1a-c) demonstrated a clear reduction in ventilation across the whole lung at 3-weeks post-tumor induction, with more subtle alterations observed at the 2-week time point. Considering that tumor burden accounted for only 1.5% of the lung area in the 2-week group, the fact that measurable changes in ventilation suggests XV could have high sensitivity in this application.

Interestingly, the severity of ventilation defects varied between animals. For example, one 3-week tumor-bearing mouse (C3) showed widespread ventilation defects, while another (C1) only exhibited mild changes. A similar pattern was also observed in the 2-week group, where one animal (B1) appeared to be more mildly affected, displaying a normal ventilation map comparable to the control animals. This variability was reflected in the global metrics and in the tumor burden quantification. For example, C3 had a markedly reduced mean specific ventilation (MSV) and tidal volume (V_T_) whereas C1 exhibited relatively normal mean specific ventilation (MSV) and normalised ventilation defect percentage (nVDP), but did show a reduction in ventilation heterogeneity (VH). The only XV parameter that was significantly altered was the large-scale ventilation heterogeneity (VH_LS_), specifically between the 3-week group and controls, however this was likely due to the small sample size and inter-animal variability.

Overall, tumor-bearing lungs tended to progress towards more homogeneous ventilation (reduced ventilation heterogeneity, VH), coupled with overall lower airflow (reduced mean specific ventilation (MSV) and increased normalised ventilation defect percentage (nVDP)), suggesting that overall the tissue is stiffer. This matches the flexiVent findings where the respiratory system (C_rs_) and static compliance (C_st_) were significantly reduced at 3-weeks. However, it is likely that as the tumors progress ventilation heterogeneity increases locally due to the patchy tumor tissue. For example, the 2-week animals in which we identified small areas of low ventilation (B2 and B3) had high small-scale ventilation heterogeneity (VH_SS_) values, which likely reflects this phenomenon. Over time the lung motion overall becomes more homogeneous, resulting in reduced large-scale ventilation heterogeneity (VH_LS_) at 3-weeks. In obstructive lung diseases, low ventilation heterogeneity (VH) indicates more homogeneous ventilation, which is typically associated with less patchy disease and preserved lung function [12]. Here the reduced ventilation heterogeneity (VH) may still reflect homogeneous ventilation, but the overall level of ventilation was reduced (i.e. low mean specific ventilation (MSV)), indicating impaired lung function. Thus, both of these parameters must be considered together to interpret lung health.

The case study of the region of interest in mouse B3 (Figure 2) adds to these findings. The whole lung ventilation, indicated by the grey line, demonstrated a typical nonlinear inflation pattern, with rapid expansion occurring early in the breath, followed by a slower increase approaching peak inspiration. In contrast, the region of interest (red line) showed an abnormal trajectory, rather than increasing in early inhalation, specific ventilation actually decreased between Phase 1 and 2. In obstructive diseases [12] or models of obstruction [11] this altered specific ventilation trajectory would likely reflect mucus obstruction resulting in delayed filling. However, in this model of lung cancer we propose that this is the result of altered tissue properties (i.e. increased stiffness) caused by tumor growth that prevents normal lung tissue expansion. Interestingly, in this 2-week animal (B3) there were not substantial differences in tissue compliance, indeed changes in compliance (C_rs_ and C_st_) did not become statistically significant until the 3-week timepoint. These findings suggest that XV imaging may be sensitive enough to detect these subtle local functional impairments not evident in the global flexiVent lung mechanics testing, or the structural CT imaging.

The flexiVent results indicated a significant decline in lung function at 3-weeks, but not at 2-weeks. At the 3-week timepoint, inspiratory capacity (IC) was significantly reduced, suggesting that the presence of tumors in the lung reduced the overall volume of air the mice were able to inhale. There was also a significant increase in overall airway resistance (R_rs_) and a decrease in compliance (C_rs_), indicating that tumors also impair the ability of the lungs to stretch and allow airflow during breathing. Pressure-volume loops further supported these findings, showing a reduction in static compliance (C_st_) and a smaller overall area under the curve, further indicating stiffer lungs with reduced inspiratory capacity (IC). The NPFE maneuver revealed a significant decrease in forced vital capacity (FVC), consistent with reduced lung volume. Other parameters (e.g. R_n_, G, FEV 0.05) were not statistically different from control, but as for the XV parameters, this was likely due to the small sample size.

The histological findings confirmed the XV and flexiVent results, with tumor burden substantially increasing between 2-weeks and 3-weeks. Interestingly, we found strong correlations between the log_10_(tumor count) and the forced expiratory volume (FEV0.05), large-scale ventilation heterogeneity (VH_LS_) and mean CT gray value, but not with any of the other parameters.

### Limitations and outlook

The primary limitation of the model is the small sample sizes that we utilised in our pilot design. We expect that with larger sample sizes many of the parameter changes would become statistically significant, potentially at the 2-week timepoint. Importantly however, we were still able to draw strong conclusions about the capabilities of XV imaging, and establish the expected changes in flexiVent mechanics in this pilot study.

As this was a proof-of-concept pilot study, feature matching between the histology, CT and XV images was not performed. As discussed by Nolte et al. (2022), this is a challenging procedure, and something that could be investigated in future studies [20].

While this mouse model recapitulates the characteristic of the human tumours at the molecular and cellular level, human tumours are predominantly one single tumor foci in a lung lobe, instead of the many lung tumor foci observed in all the lobes in this mouse model (caused by systemic delivery of the cancer cells). The large overall tumor burden in the mice would impact lung function in the whole volume of the lung, while in the human, tumors would affect a more restricted region of the lung. This suggests there is likely to be significant value from moving toward human clinical trials rather than performing larger studies in mouse models. Nonetheless, that approach also presents challenges as human lung cancer patients often have comorbidities such as chronic obstructive pulmonary disease or fibrosis that complicate assessments.

Future longitudinal rodent studies could be used to track tumor progression in individual animals, and enable improved evaluation of chemotherapeutics. The XV and CT scans are not necessarily terminal procedures. In this study animals were only euthanised because a tracheostomy was required for successfully performing the selected flexiVent measurements. As we showed here, there can be significant differences in tumour growth rates, so longitudinal tracking of individual animals could reduce the total number of animals needed for studies and improve statistical power. However, the effect of any radiation from the repeated scanning on tumours would need to be understood.

As with all papers using XV imaging, this study refers to the data produced by XV as measuring ventilation, however what is being measured is actually tissue expansion. In obstructive lung diseases, the elasticity of the tissue remains approximately constant while the force being applied by air is reduced due to obstructed airways. This leads to a true reduction in ventilation, which is visible due to the lung reduced expansion of the lung tissue. However in this model, the elasticity of the tissue has changed, and so tissue expansion and true ventilation are not analogous, so further studies are required to fully understand how tissue properties such as stiffness affect the XV data, and how this is altered by tumor growth.

### Clinical significance

In patients, tumour burden and response to therapy are primarily monitored by anatomical imaging (e.g. CT, MRI) and nuclear medicine (e.g. PET). These primarily assess either the tumour size or metabolic activity, not its direct impact on lung function. These preliminary results suggest combining anatomical imaging with quantitative, region-specific XV functional imaging could: (i) provide earlier or more sensitive detection of clinically significant tumour progression; (ii) help distinguish between stable disease on imaging but declining lung function (or vice versa); (iii) support decision-making for therapies or interventions (e.g. surgery vs. continued chemotherapy).

Early functional changes detectable by XV may precede detectable anatomical abnormalities. Therefore, identifying subtle shifts in ventilation before significant structural changes appear on imaging could allow clinicians to identify patients at risk of rapid decline much earlier. If functional impairment – rather than just tumor size – is predictive of outcomes or complications such as respiratory failure, infection, or tolerance to surgery, then XV functional imaging could be used to stratify patients for treatment risk or need for supportive care. XV ventilation imaging may provide value for guiding radiotherapy or surgical planning. Regionally specific impairment – as noted in our case study of mouse B3 – may guide surgeons or radiation oncologists to target or spare particular lung segments that are responsible for significant ventilation (e.g. Australian clinical trial ACTRN12612000775819).

## Conclusion

Using novel non-invasive *in vivo* XV lung function imaging in conjunction with gold-standard flexiVent whole-lung mechanics, we have performed a detailed study of lung function in a mouse model of lung cancer; the first of its kind. The results suggest that the severity of the model increases dramatically between weeks 2 and 3, with both the flexiVent and XV imaging detecting statistically significant changes between the control and 3-week groups, but not the 2-week group. Future human XV clinical trials, similar to those being performed for cystic fibrosis and other lung diseases, could reveal more about the alterations to airflow from lung cancer.

## Acknowledgements

The authors acknowledge the facilities and scientific and technical assistance of the National Imaging Facility, a National Collaborative Research Infrastructure Strategy (NCRIS) capability, at the Preclinical Imaging and Research Laboratories, South Australian Health and Medical Research Institute. The lung cancer cells were a kind gift from Dr K. Sutherland.

## CRediT Contributions

RS: Data Curation, Formal analysis, Investigation, Software, Visualization, Writing - Original Draft

NR: Formal analysis, Investigation, Visualization, Writing - Original Draft

DB: Data Curation, Formal analysis, Investigation, Visualization, Writing – Review & Editing

NE: Investigation, Methodology, Writing - Original Draft

MLAL: Conceptualisation, Resources, Funding acquisition, Methodology, Writing - Original Draft

MD: Conceptualisation, Data Curation, Formal analysis, Funding acquisition, Investigation, Methodology, Supervision, Visualization, Writing - Original Draft

## Data Availability

The data generated in this study is available at DOI: 10.25909/29312042

